# Transparent and stretchable metal nanowire composite recording microelectrode arrays

**DOI:** 10.1101/2022.10.11.511842

**Authors:** Zhiyuan Chen, Khanh Nguyen, Grant Kowalik, Xinyu Shi, Jinbi Tian, Mitansh Doshi, Bridget R. Alber, Xin Ning, Matthew W. Kay, Luyao Lu

## Abstract

Transparent microelectrodes have received much attention from the biomedical community due to their unique advantages in concurrent crosstalk-free electrical and optical interrogation of cell/tissue activity. Despite recent progress in constructing transparent microelectrodes, a major challenge is to simultaneously achieve desirable mechanical stretchability, optical transparency, electrochemical performance, and chemical stability for high-fidelity, conformal, and stable interfacing with soft tissue/organ systems. To address this challenge, we have designed microelectrode arrays (MEAs) with gold coated silver nanowires (Au-Ag NWs) by combining technical advances in materials, fabrication, and mechanics. The Au coating improves both the chemical stability and electrochemical impedance of the Au-Ag NWs microelectrodes with only slight changes in optical properties. The MEAs exhibit a high optical transparency >80% at 550 nm, a low normalized 1 kHz electrochemical impedance of 1.2-7.5 Ω cm^2^, stable chemical and electromechanical performance after exposure to oxygen plasma for 5 minutes and cyclic stretching for 600 cycles at 20% strain, superior to other transparent microelectrode alternatives. The MEAs easily conform to curvilinear heart surfaces for co-localized electrophysiological and optical mapping of cardiac function. This work demonstrates that stretchable transparent metal nanowire MEAs are promising candidates for diverse biomedical science and engineering applications, particularly under mechanically dynamic conditions.

## 1. Introduction

Recording cellular activity at high spatiotemporal resolution from soft tissue/organ systems with microelectrodes is a critical component of many biomedical research and clinical applications, including basic physiology studies, investigations of disease, and the development of disease therapies.^[1-4]^ Despite their tremendous impact on classic electrophysiological research, microelectrodes are unable to readout important functional parameters such as intracellular calcium dynamics, metabolic activity, or target specific cell types. In recent years, optical techniques such as high speed fluorescence imaging have been widely used to measure important cellular parameters (e.g., membrane potentials and intracellular calcium) and optogenetics has been used to manipulate cells or circuits with cell-type specificity through the incorporation of light-sensitive proteins.^[5-8]^ Insights gained from such optical sensing and actuation is valuable and complimentary to information provided by microelectrode recordings. In this context, combining electrical and optical modalities will yield competentary tools that leverage the advantages of both techniques. However, conventional electrophysiological studies rely heavily on opaque metal (e.g., gold, platinum, and iridium) microelectrodes, which unfortunately produce severe photoelectric artifacts that distort the recorded signals and prevent direct optical probing of cells at the same sites.^[9-11]^

Recent developments in optically transparent microelectrodes such as indium tin oxide (ITO),^[12]^ carbon-based materials,^[13-14]^ and metal nanostructures,^[15-17]^ have endowed efficient light delivery through the microelectrodes with negligible photoelectric artifacts.^[17]^ In addition to optical transparency, sufficient mechanical stretchability is required to achieve conformal contact with soft, curvilinear tissue surfaces and withstand the repetitive mechanical strains induced by the tissue/organ during vital body functions such as heart contraction, breathing, and walking (e.g., strain levels can exceed 10% in the brain and 20% in the heart).^[18-20]^ The large mechanical mismatch between conventional rigid microelectrodes and soft tissue is the cause of many issues, such as tissue damage, inflammation, large stress at the device interfaces, delamination of microelectrodes from tissue surfaces, and microelectrode performance degradation over time,^[21-23]^ making effective electrophysiological recording from the surrounding tissue challenging and limiting the success of current microelectrode designs. Among commonly used transparent microelectrode materials, ITO is unsuitable for stretchable applications due to its brittle nature.^[24]^ While graphene presents remarkable optical properties and chemical stability, the carbon-carbon network of graphene is easily broken, which limits the stretchability of monolayer graphene to <5% due to a lack of strain dissipation.^[25-26]^ In addition, both graphene and carbon nanotubes (CNTs) suffer from a complex fabrication process and undesirable high electrochemical impedance. Silver nanowires (Ag NWs) are a low-cost stretchable transparent electrode material that are widely used in traditional optoelectronic devices. Ag NWs are easy to fabricate on a large scale and have excellent electrical, optical, and mechanical properties.^[27-29]^ Solution-processed Ag NWs networks also exhibit inherent high surface roughness because they are not strictly in the same film plane, which increase the effective interfacial contact area between the microelectrodes and the electrolytes, markedly boosting the electrochemical performance for electrophysiological recording applications.^[30-32]^ Although attractive, direct exposure of Ag to tissue could have adverse health effects.^[33]^ Ag NWs can also be highly corrosive in biofluids due to oxidation, which reduces their long-term stability.^[34-35]^ These critical issues must be addressed to realize the full potential of the metal NWs-based stretchable and transparent microelectrode technologies.

To overcome these issues, we developed and tested optically transparent, mechanically stretchable, and chemically stable microelectrode arrays (MEAs) constructed from patterned gold-coated Ag NWs (Au-Ag NWs) composites on thin elastomer substrates. The MEAs exploit advances in materials, mechanics, and micro-/nano-fabrication schemes, which serve as foundations to address the current challenges in developing the next generation of high-performance soft transparent MEA technologies that can maintain contact and mechanically conform to curved tissue surfaces. The ultrathin conformal Au coating greatly improves the electrochemical performance and oxidation resistance of the Au-Ag NWs without sacrificing optical transparency. The resulting Au-Ag NWs microelectrodes exhibit high optical transparency >80% at 550 nm, low normalized 1 kHz electrochemical impedance of 1.2-7.5 Ω cm^2^, superior oxidation resistance with negligible changes in electrochemical performance after direct exposure to oxygen plasma for 5 minutes, and excellent mechanical stretchability and durability (up to 600 cycles at 20% tensile strain). The Au-Ag NWs fabrication process provides MEAs with high performance parameters that are consistent between MEAs. To test the MEAs in a mechanically active biological environment, the Au-Ag NWs MEAs were applied to the epicardium of excised perfused rat hearts where electrical activity was mapped from the same area simultaneously using the MEAs and via fluorescence imaging of a potentiometric dye (optical mapping). Local tissue activation times and conduction velocities during normal sinus rhythm and electrical pacing were the same when measured using the MEAs or optical mapping, demonstrating faithful co-localized recording of cardiac activity with the MEAs and optical mapping through the MEAs. In summary, the techniques presented here to integrate novel nanomaterials, stretchable electronics, and transparent microelectrodes into a single device will inspire the development of human-machine interfaces with sophisticated bioelectronic systems to enable a broad spectrum of applications in multimodal cellular/tissue mapping and modulation.

## 2. Results and Discussion

### 2.1. Design and Fabrication of Au-Ag NWs MEAs

Figure 1 presents a schematic illustration of the fabrication process of Au-Ag NWs MEAs. First, a poly(methyl methacrylate) (PMMA) sacrificial layer is coated on a handling glass. A 7 μm transparent and biocompatible SU-8 adhesive layer is spin-coated on PMMA, followed by spin-coating of Ag NWs/isopropyl alcohol (IPA) solutions. Photolithography generates Ag NWs/SU-8 structures into the shape of the predesigned serpentine connected 9-channel MEAs with Ag NWs partially embedded in SU-8 to prevent nanowire delamination under mechanical deformations.^[31]^ The serpentine structures enable excellent mechanical properties of the MEA. The pitch value (center to center distance between two closest microelectrodes in the MEA) is 2 mm, similar to the space constant of cardiac tissue.^[36]^ Another 7 μm SU-8 epoxy encapsulates the Ag NWs and defines the microelectrode areas. An ultrathin Au layer (6 nm) is conformally coated on the surfaces of Ag NWs in the microelectrode areas using electroplating. A water-soluble tape (WST) picks up the Au-Ag NWs MEA patterns after the sacrificial PMMA layer dissolves in acetone. An ultrathin (50 nm) transparent SiO_2_ is deposited on the backside of the MEAs. The MEAs attached WST and a 35 μm-thick ultrasoft, biocompatible, transparent polydimethylsiloxane (PDMS) elastomer substrate are exposed to oxygen plasma treatment, attached together and heated to create strong Si-O-Si chemical bonding between the MEAs and PDMS. Removing the WST completes the fabrication. The total thickness of the MEAs is ∼50 μm. Notably, the solution processing method to pattern the MEAs is scalable, and the number of microelectrodes and pitch values can be scaled up or down based on specific research needs. The fabrication details are summarized in the Experimental Section.

**Figure 1.**
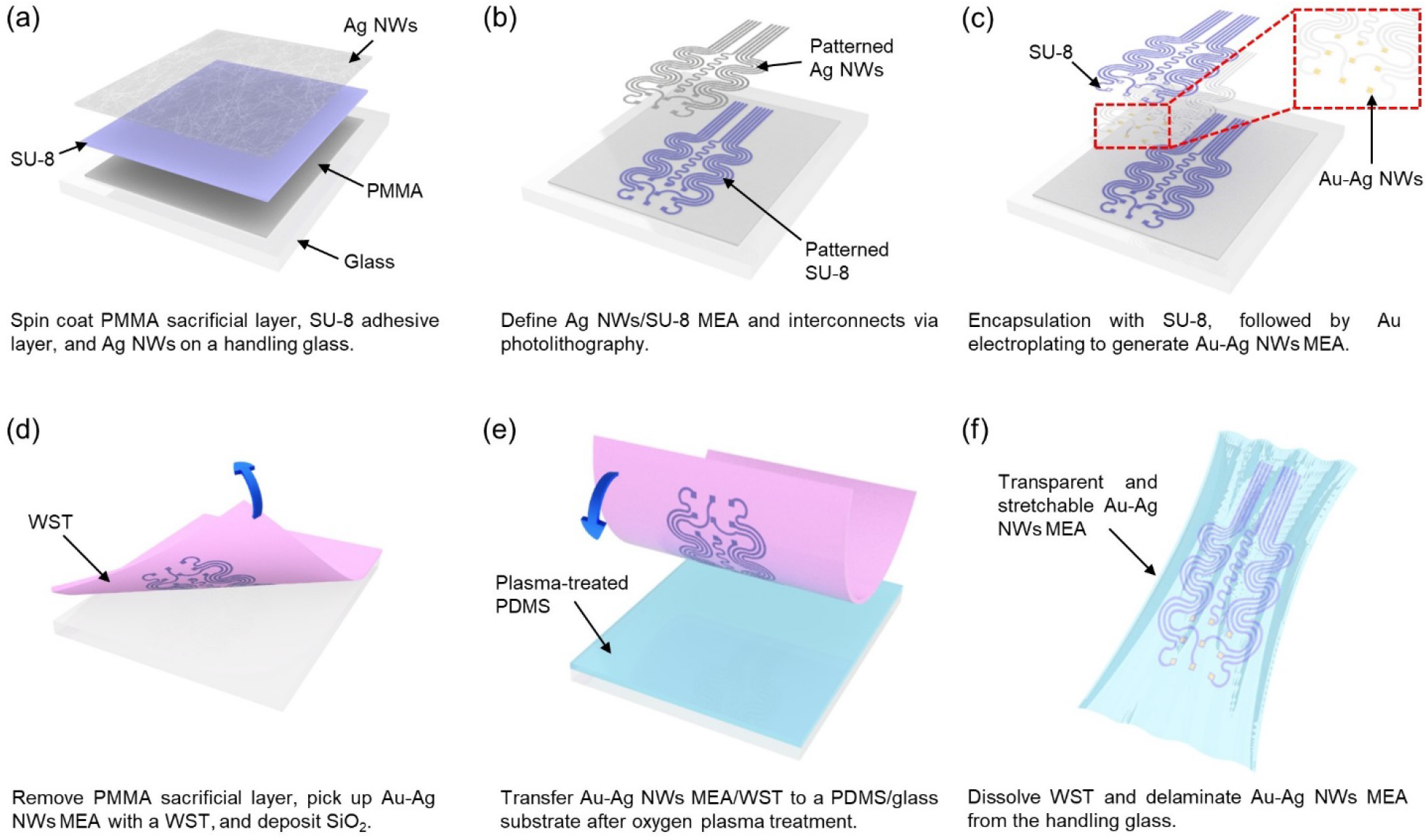
Schematic illustration of detailed steps for fabricating the transparent and stretchable Au-Ag NWs MEAs.

### 2.2. Optical and Electrical Properties of Au-Ag NWs Structures

**Figure 2**a presents an optical image of a highly transparent 9-channel Au-Ag NWs MEA device (optical transmittance ∼80% at 550 nm). Scanning electron microscopy (SEM) image in Figure 2b reveals the fine Au-Ag NWs networks in the microelectrode areas. The NWs have a diameter of ∼100 nm and a length of ∼25-40 μm. The uniform and conformal Au coating (6 ± 0.2 nm) and core-shell structure of the Au-Ag NWs is confirmed by transmission electron microscopy (TEM) results in Figure 2c. The nanowire network density is a critical design parameter in determining the resulting optical, electrical, and electrochemical properties of the microelectrodes. Here, the network density is controlled by the concentrations of Ag NWs in IPA. Figure 2d presents the transmission spectra of Ag NWs networks in the range of visible wavelength from 400 to 700 nm. As expected, the average transmittance values at 550 nm decrease from 81.6 ± 1.05% to 70.6 ± 1.33%, and 60.8 ± 1.84% with the Ag NWs concentrations increasing from 3 to 5, and finally 10 mg/mL, respectively. After electroplating the Au shell layer, the average transmittance values exhibit only a slight decrease (∼1-2%) and change to 80.1 ± 2.15%, 69.8 ± 1.86%, and 60.0 ± 2.45%, respectively (Figure 2e). This is because the diameter of the pristine Ag NWs is much larger than the Au shell layer. It it well-known that Au coating reduces the wire-to-wire junction resistance of Ag NWs.^[37]^ The sheet resistance values of the Ag NWs interconnects show a moderate decrease from 15 and 20 Ω sq^-1^ at high transmittance values of 70.6% and 81.6% (Figure 2f) to 12 and 17 Ω sq^-1^ after electroplating the Au layer. Those results outperform that of CNTs (150-500 Ω sq^-1^),^[18]^ graphene (76-152 Ω sq^-1^),^[13]^ and ITO polyethylene terephthalate films (60 Ω sq^-1^)^[13]^ at similar transmittance levels. As a result, the nanowire networks can simultaneously serve as efficient transparent interconnects for the MEAs to simplify the overall device fabrication process.

**Figure 2.**
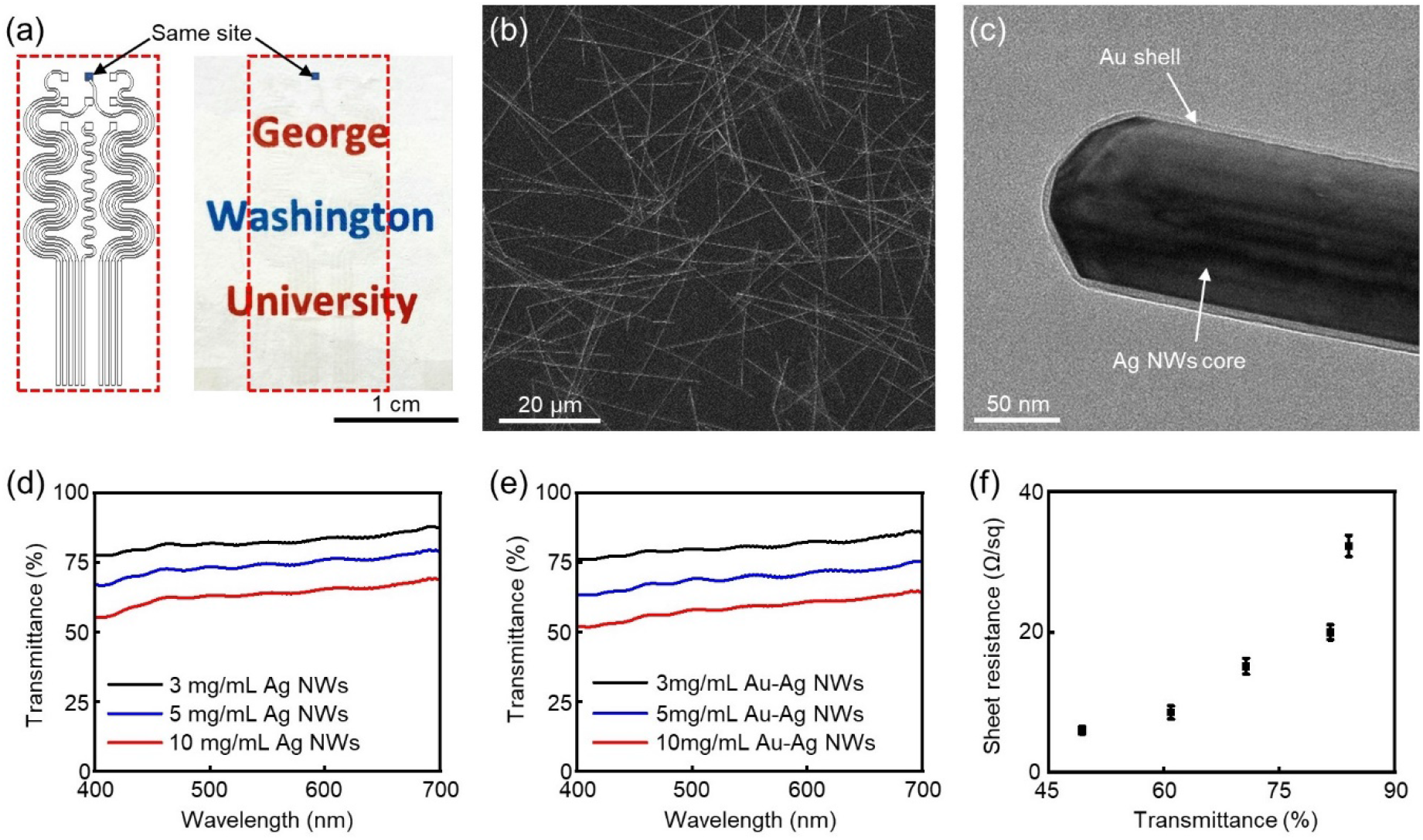
a) (Left) Schematic illustration of the MEA and interconnects layout. (Right) Optical image of a highly transparent, stretchable, 3 × 3 Au-Ag NWs MEA and interconnects. The Ag NWs concentration is 3 mg/mL. The MEA and interconnects area is marked by the red dashed box. b) SEM image of the Au-Ag NWs networks in the microelectrode in (a). c) TEM image of a core-shell structured Au-Ag NW. Transmission spectra of Ag NWs at different concentrations (3, 5, and 10 mg/mL) without (d) and with (e) Au coating. f) Sheet resistance of the Ag NWs interconnects versus optical transparency.

### 2.3. Electrochemical Properties of Au-Ag NWs Microelectrodes

A low electrochemical impedance is desired to achieve high signal-to-noise ratios (SNRs) for efficient electrophysiological recording. The electrochemical impedance is measured using electrochemical impedance spectroscopy (EIS) at frequencies from 1 Hz to 100 kHz in a phosphate-buffered saline (PBS). This frequency band includes the frequencies associated with extracellular and intracellular potential signals. **Figure 3**a presents the electrochemical impedance of pristine Ag NWs microelectrodes as a function of concentrations. The 1 kHz electrochemical impedance values increase from 3.2, to 11, and 26 kΩ with Ag NWs concentrations decreasing from 10, to 5, and 3 mg/mL, respectively. Electroplating the Au layer significantly improves the electrochemical performance with the 1 kHz electrochemical impedance values decrease to 8.6 (3 mg/mL), 3.9 (5 mg/mL), and 0.80 (10 mg/mL) kΩ, respectively (Figure 3b). We propose that the improved electrochemical impedance results from the combined effects of improved conductivity, increased surface area, and the improved electrochemical performance of Au compared to Ag. Figure 3c illustrates the electrochemical impedance spectra of solid Au and Ag film microelectrodes (300 × 300 μm^2^ and 100 nm thick), where Au microelectrodes show lower electrochemical impedance values than Ag microelectrodes across the entire frequency range. The 1 kHz electrochemical impedance of the Au-Ag NWs microelectrodes at 10 mg/mL Ag NWs processing condition (0.80 kΩ) is on par with that of the solid Au film microelectrode (2.3 kΩ). The electrochemical and optical performance of Au-Ag NWs microelectrodes are further compared to other transparent microelectrode counterparts, including CNT,^[18]^ graphene,^[14]^ ITO,^[38]^ and metal nanostructures^[16, 24, 39]^ (Figure 3d). The Au-Ag NWs microelectrodes exhibit a superior normalized electrochemical impedance (7.5 Ω cm^2^ at 1 kHz) at a high transmittance (80.1%), which are among the most competitive performance for soft transparent microelectrodes used in electrophysiological recording studies. The phase response of Au-Ag NWs microelectrodes shows they are more capacitive (phase angles between -85° and -60°) at low frequencies (1 Hz to 1 kHz) and become more resistive at high frequencies (Figure 3e). In view of the balanced electrochemical and optical properties, Au-Ag NWs microelectrodes with the normalized electrochemical impedance of 7.5 Ω cm^2^ and transmittance of 80.1% are used in subsequent experiments. Figure 3f shows the highly uniform electrochemical impedance responses from all 9 microelectrodes in a 3 × 3 Au-Ag NWs MEA with an average 1 kHz electrochemical impedance value of 8.3 ± 1.4 kΩ, respectively, which ensures high-fidelity electrophysiological mapping across different sites. Our fabrication method enables the patterning of microelectrodes down to a cellular scale. Figure S1, Supporting Information presents the electrochemical impedance spectrum of a Au-Ag NWs microelectrode with a dimension at 50 × 50 μm^2^, comparable to the sizes of individual cardiomyocytes.^[40]^ The microelectrode exhibits a moderate 1 kHz electrochemical impedance of 237 kΩ at such a small size.

**Figure 3.**
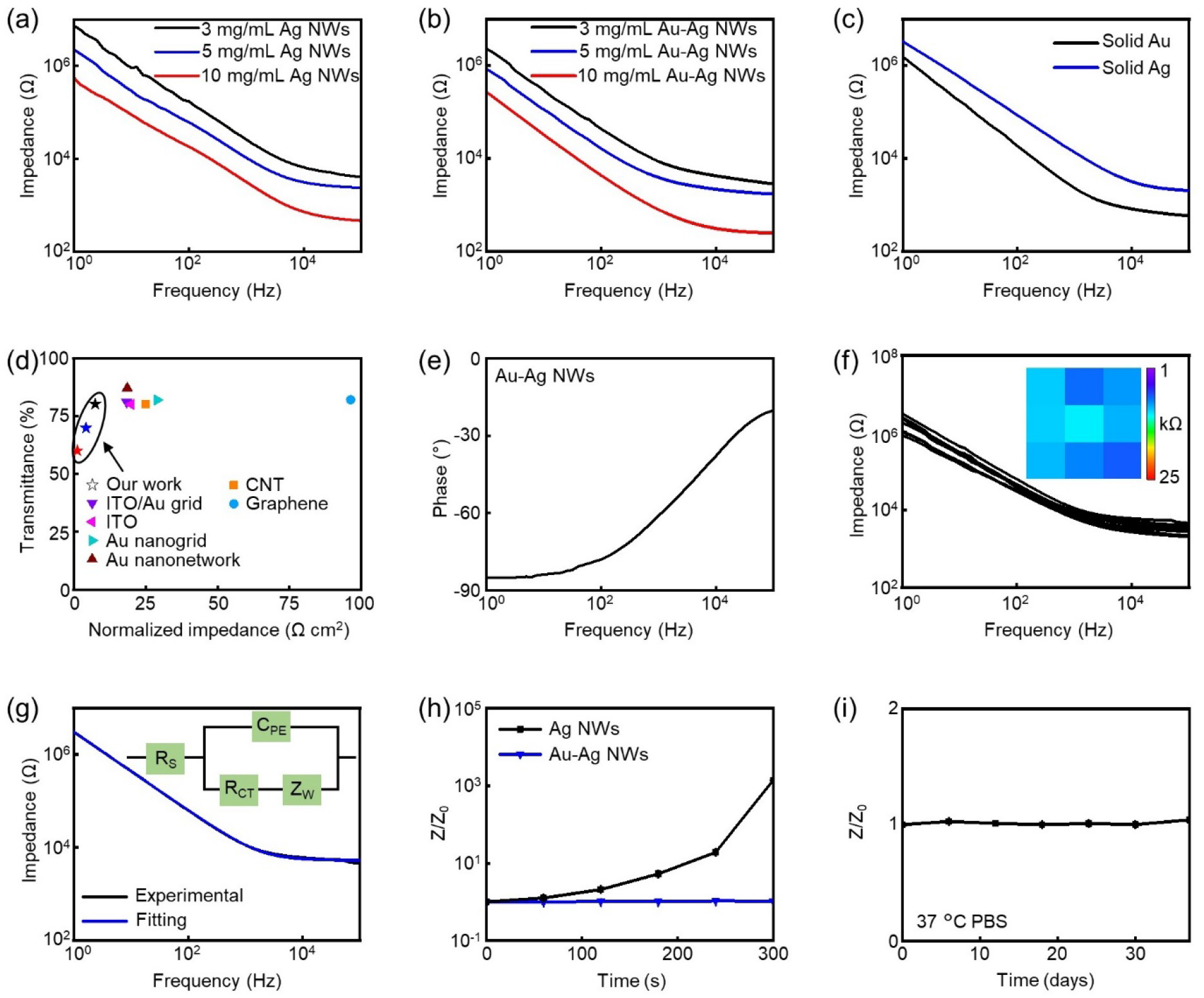
Impedance plots of Ag NWs microelectrodes at different Ag NWs concentrations (3, 5, and 10 mg/mL) without (a) and with (b) Au coating. The size of the microelectrodes is 300 × 300 μm^2^. c) Impedance plots of 300 × 300 μm^2^ opaque solid Au and Ag film microelectrodes. d) Comparison of normalized impedance and transmittance of the Au-Ag NWs microelectrodes to major reported transparent microelectrodes. e) Phase plot of the Au-Ag NWs microelectrodes. f) Impedance plots of all 9 microelectrodes in a 9-channel Au-Ag NWs MEA. Inset: 1 kHz impedance colormap with respect to actual microelectrode position in the MEA. g) Equivalent circuit model fitting of the Au-Ag NWs microelectrodes. h) Electrochemical performance stability of Ag NWs and Au-Ag NWs microelectrodes under oxygen plasma treatment. Z_0_ is the initial impedance, whereas Z represents the impedance at a specific oxygen plasma exposure time. i) Soak test of the Au-Ag NWs microelectrodes after immersed in a PBS solution at 37 °C for 5 weeks. Z_0_ is the initial impedance, whereas Z represents the impedance on a specific day.

The EIS results of Au-Ag NWs microelectrodes are fit to an equivalent circuit model in Figure 3g (inset) to gain more insights into the microelectrode-electrolyte interfacial properties. The elements in the circuit model include solution resistance (R_S_), charge transfer resistance (R_CT_), Warburg element for diffusion (Z_W_), and parallel constant phase element (C_PE_).^[31]^ The experimental results match well with the fitting results (Figure 3g). C_PE_ is defined by 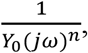 where *Y*_*0*_ is the magnitude of C_PE_, *j* is the unit imaginary number, *ω* is the angular frequency, and *n* is a constant (0 ≤ n ≤ 1). When n = 0 it represents a pure resistor while when n = 1 it represents an ideal capacitor. **Table 1** summarizes the fitting parameters. The n value (0.85) for the Au-Ag NWs microelectrodes represents a more double-layer capacitive interface, which is desired for electrophysiological recording applications.

**Table 1.** Equivalent circuit fitting results of Au-Ag NWs microelectrodes.

| $R_S (\Omega)$ | $C_{PE} (S \times s^n)$ | | $R_{CT} (\Omega)$ | $Z_W (S \times s^{1/2})$ |
| --- | --- | --- | --- | --- |
|  | Q | n |  |  |
| $5.3 \times 10^3$ | $7.0 \times 10^{-8}$ | 0.85 | $1.5 \times 10^6$ | $4.3 \times 10^{-13}$ |

### 2.4. Chemical Stability of Au-Ag NWs Microelectrodes

Ag NWs are easily oxidized when exposed to body fluids, resulting in decreased performance over time and releasing of Ag ions into the biological environment.^[41-42]^ The chemical stability of the microelectrodes is significantly improved by the electroplated Au shell layer since Au is an inert noble metal with a higher reduction potential than Ag. When the bare Ag NWs are oxidized, the formation of silver oxides on the surface of the Ag NWs will significantly increase the electrochemical impedance. The oxidation resistance of the Au-Ag NWs is evaluated using oxygen plasma as the oxidant. The Au-Ag NWs microelectrodes show no changes in electrochemical impedance after direct exposure to oxygen plasma for 5 minutes while the electrochemical impedance of the bare Ag NWs microelectrodes increases >1,300 times under the same condition (Figure 3h). Those results indicate the successful protection of Ag NWs from oxidation by the Au shell layer and strong adhesion between the Au shell and Ag NWs core. The long-term stability of the Au-Ag NWs microelectrodes is assessed by monitoring their electrochemical impedance for 5 weeks with the microelectrodes placed in a 37 °C PBS solution. Negligible performance changes are observed, suggesting no delamination or corrosion of the nanowires over time in the aqueous solutions.

### 2.5. Mechanical Properties of Au-Ag NWs Microelectrodes

**Figure 4**a shows that the Au-Ag NWs microelectrodes exhibit stable electrochemical performance during one-time stretching measurements with up to 40% tensile strain, defined as the relative length increase applied to the MEAs. Cyclic stretch and release measurements at 20% tensile strain evaluate the mechanical stability and durability of the Au-Ag NWs MEAs. No changes in the electrochemical impedance are observed after 600 stretch and release cycles (Figure 4b). Simulations by finite element analysis (FEA) show the strain distribution in the Au-Ag NWs MEAs under different strain levels, from 0% to 40%, (Figure 4c and Figure S2, Supporting Information). All device layers are perfectly bonded in FEA and the center serpentine region experiences higher strain as compared to other serpentine regions in the structure. The electromechanical stability and electrophysiological recording capability are further demonstrated by recording a programmed sine wave signal (10 Hz, 20 mV peak-to-peak amplitude) in a PBS solution using the microelectrodes before and after 500 cycles of stretching at 20% tensile strain. Figure 4d shows that the recorded sine waves exhibit no observable decreases in signal amplitude, suggesting the robust mechanical performance of the microelectrodes. Figure S3 (Supporting Information) presents the power spectrum density (PSD) of the benchtop recording signals to provide details on noise and signal in the frequency domain. The large peaks at 10 Hz are from the input sine wave signal. The Au-Ag NWs MEAs show uniform and stable SNRs from all 9 channels before (44.8 ± 0.3 dB) and after (44.2 ± 1.0 dB) 500 cycles of stretch and release at 20% tensile strain (Figure 4e). Collectively, these experimental and FEA results highlight the robust elastic responses and mechanical compliance of the stretchable Au-Ag NWs MEAs at strain levels relevant to realistic physiological conditions.

**Figure 4.**
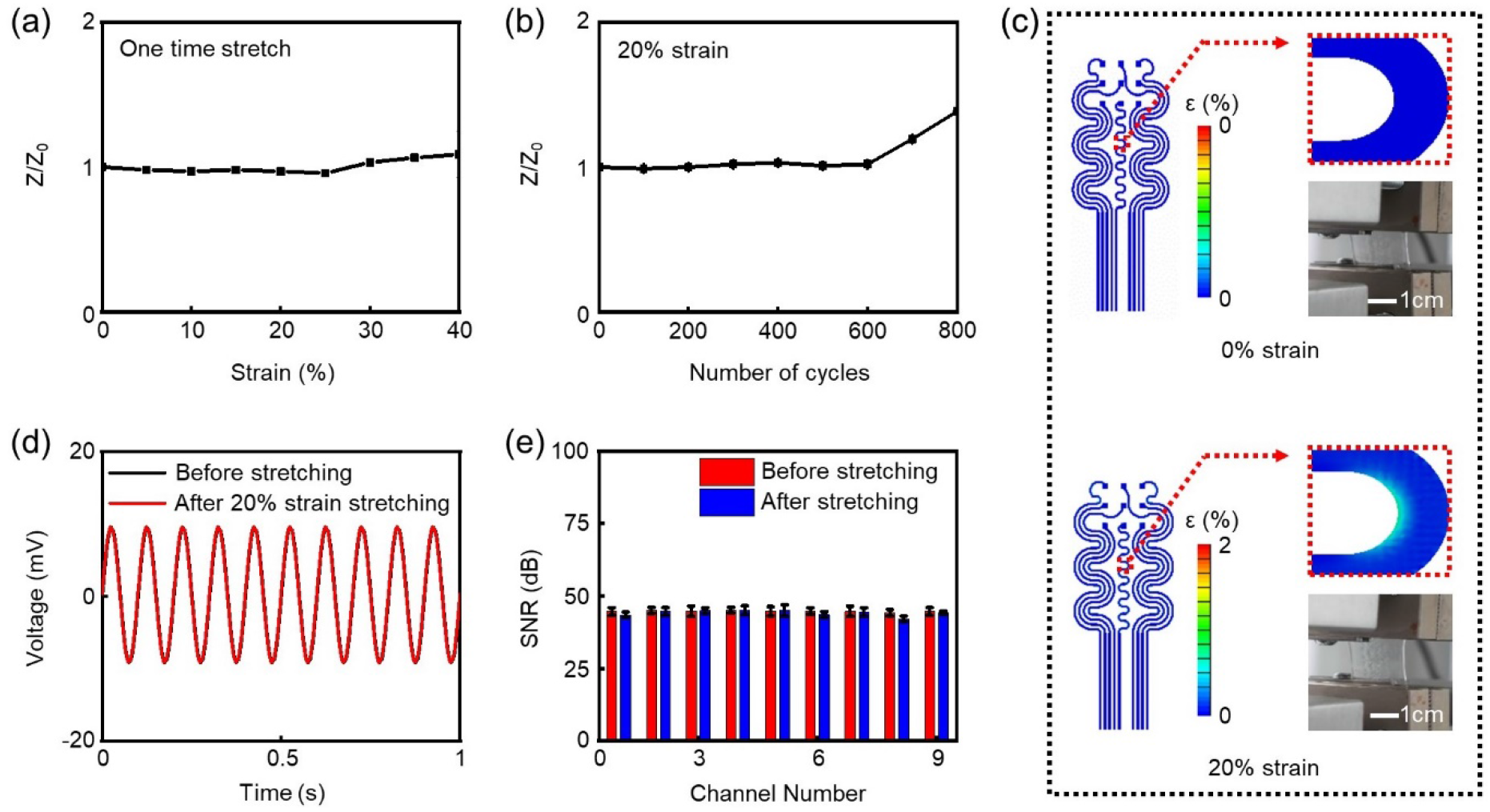
Variation of electrochemical impedance after one-time stretching (a) and continuous stretch and release at 20% strain (b) for the Au-Ag NWs MEAs. Z_0_ is the initial impedance, whereas Z represents the impedance at a specific strain or stretching cycle. c) FEA strain distribution results and optical images of the Au-Ag NWs MEAs under 0% and 20% uniaxial stretching. d) Benchtop recording results from the pristine and stretched (20% strain for 500 cycles) Au-Ag NWs microelectrodes, with 20 mV peak-to-peak amplitude and 10 Hz sine wave input. e) SNR histogram of the 9 microelectrodes in a pristine and stretched Au-Ag NWs MEA from the benchtop electrical recording in (d).

### 2.6. Co-localized Cardiac Electrical and Optical Mapping Using Au-Ag NWs MEAs

MEAs that intimately interface with distinct regions of the heart for monitoring the spatiotemporal dynamics of cardiac activity are crucial for basic cardiac studies and therapeutic applications. Here, *ex vivo* experiments using excised perfused adult rat hearts were conducted to validate the Au-Ag NWs MEAs for co-localized crosstalk-free spatiotemporal electrical and optical measurements. **Figure 5**a (left) illustrates the experimental setup for electrical mapping and pacing, including the three electrocardiogram (ECG) electrodes in the superfusate bath and the epicardial location of the 9 microelectrodes in the Au-Ag NWs MEAs. After being stretched for 500 cycles at 20% tensile strain, the stretchable and transparent MEA conformally contacts the left ventricle epicardium, as shown in Figure 5a (right). Figure 5b shows the time-aligned ECG recorded from the bath electrodes and the 9 electrogram (EG) signals recorded from the MEA during sinus rhythm. High frequency spikes in each EG signal align with the R wave of the ECG, indicating synchronized local depolarization of tissue beneath each Au-Ag NWs microelectrode. The average heart rate is 237 ± 0.26 beats per minute (BPM) from the MEA EG results, which matches well with that (237 ± 0.30 BPM) of the ECG. Figure 5c displays optical and MEA EG activation maps of the epicardium during sinus rhythm that were reconstructed from depolarization times measured from optically mapped action potentials and the MEA EGs. The strong correlation between the two maps within the region covered by the MEA EG array (black box) suggests the high-fidelity mapping capabilitiy of the Au-Ag NWs MEA. For optical mapping, the potentiometric dye RH237 with excitation and emission maxima at 520 and 782 nm, respectively, is used to transduce changes in cellular membrane potential into a fluorescence signal that can be imaged as an optical action potential.^[43]^ The acquisition of high fidelity optical action potentials through the MEA demonstrates that the high optical transmittance values of >80% of the MEA at those two wavelengths allow efficient photon delivery to and collection from the tissue below the MEA for effective co-localized electrical and optical mapping. Figure 5d displays optical and MEA EG activation maps during slow (240 BPM) and fast (545 BPM) electrical pacing from the electrode placed at the base of the heart. Again, there is strong correlation between the two maps within the region covered by the MEA EG array. Each map clearly shows propagation of cellular depolarization across the epicardium from the pacing site (above and left of EG channel 1) toward the apex, below the array. Activation maps at the two pacing rates reveal cardiac conduction velocity restitution, where the propagation speed of depolarization waves slows during fast heart rates.^[44]^ Average conduction velocity calculated from the same region that was electrically and optically mapped was 95.8 (both electrical and optical) cm s^-1^ during slow pacing and 54.4 (electrical) and 53.0 (optical) cm s^-1^ during fast pacing. Time-aligned MEA EG signals and action potentials simultaneously optically mapped from the tissue below each Au-Ag NWs microelectrode are plotted together in Figure 5e for each pacing rate. Downward deflections in the EG signals that occur after the pacing spikes align with the depolarization phase of the co-located optical action potentials, further highlighting the ability of the Au-Ag NWs MEAs to record extracellular depolarizations simultaneously with optically mapped action potentials.

**Figure 5.**
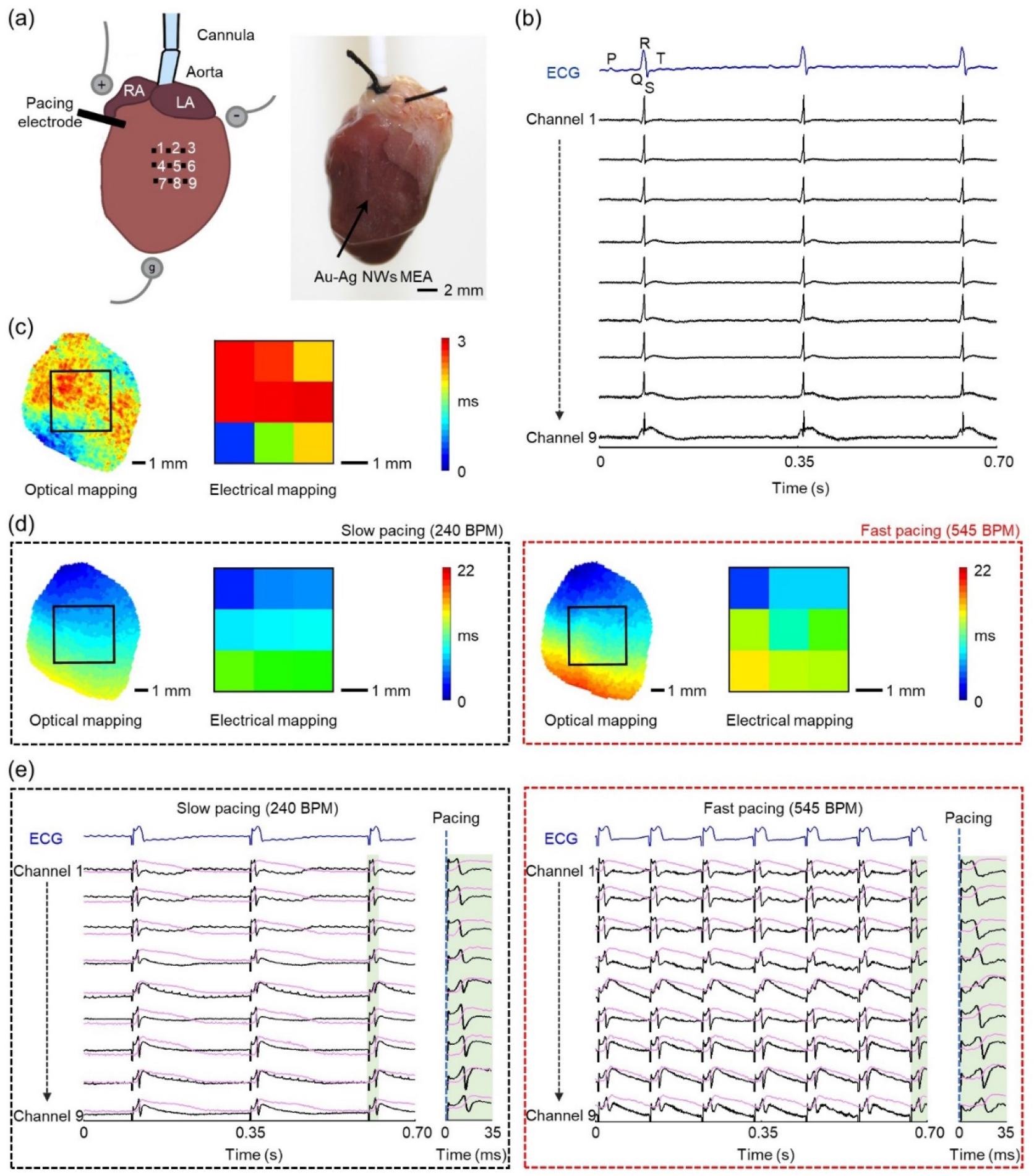
a) Left, schematic of the excised perfused rat heart experiments. Right, optical image of a stretchable, transparent Au-Ag NWs MEA conforming to the left ventricle epicardium of a perfused rat heart. b) The bath-conducted ECG and the 9 EG signals recorded from the Au-Ag NWs MEA during sinus rhythm. c) Optical (left) and electrical (right) activation maps during sinus rhythm. The black box on the optical map indicates the region mapped by the 9 MEA EGs. d) Optical and electrical activation maps during slow (left) and fast (right) electrical pacing at 240 and 545 BPM, respectively. e) Representative bath-conducted ECG signals (blue), co-localized 9-channel EG signals (black) and optical mapping signals (magenta) captured simultaneously during slow (left) and fast (right) electrical pacing conditions in (d).

## 3. Conclusion

This work demonstrates Au-Ag NWs-based stretchable and transparent MEAs that overcome the limitations of conventional rigid and opaque metal MEAs for multimodal electrophysiological and optical biointerfacing under mechanically active conditions, which represents a significant technological advancement in bioelectronics. Electroplating of an ultrathin Au layer on Ag NWs surface simultaneously improves the chemical stability and electrochemical performance without significantly sacrificing optial transparency. The resulting Au-Ag NWs microelectrodes exhibit excellent optical transparency >80%, low electrochemical impedance of 0.80-8.6 kΩ at 1 kHz, superior oxidation resistance under oxygen plasma treatment and chronic soak test in a PBS solution for >1 month, and durable mechanical performance under cyclic stretching of 600 cycles at 20% tensile strain. Successful proof-of-concept demonstrations in cardiac electrophysiology experiments demonstrate that the Au-Ag NWs MEAs enable high-fidelity co-localized extracellular electrical and action potential optical mapping of excised perfused hearts during sinus rhythm and pacing. Our results greatly expand the landscape for MEA technologies and present high-performance mechanically compliant metal nanowires-based MEAs as promising tools to interrogate soft biological systems via multiple measurement modalities.

### 4. Experimental Section

#### MEA Fabrication

A 90 nm layer of PMMA (Microchem) was spin-coated at 3000 rpm on a glass substrate. A 7 μm-thick SU-8 photoepoxy layer (SU-8 2007, Microchem) was then spin-coated on PMMA at 3000 rpm, followed by soft baking at 65 °C and 95 °C for 2 minutes each. 80 μL of Ag NWs/IPA solution (ACS Material) was spin-coated on SU-8 at 1000 rpm. Photolithography then defined the Ag NWs/SU-8 MEA and interconnect patterns. The photolithography process was carried out by exposing the Ag NWs/SU-8 mixtures to ultraviolet light with a dose of 90 mJ/cm^2^, post exposure baking at 95 °C for 4 minutes, developing in the SU-8 developer solution for 120 s, followed by a hard bake at 150 °C for 20 minutes. Another 7 μm-thick SU-8 photoepoxy layer (SU-8 2007, Microchem) defined the MEA windows *via* photolithography (spin-coating rate, 3000 rpm; dose, 90 mJ/cm^2^). A 6 nm Au layer was galvanostatically electroplated on the unencapsulated Ag NWs surface using a sulfite-based solution (TSG-250, Transene) under agitation at a current density of 0.1 mA/cm^2^ with a Gamry potentiostat (Reference 600+, Gamry Instruments Inc.). The solution was diluted by water at 1:8 volume ratio. Next, the Au-Ag NWs MEAs were immersed in acetone for 1 hour to fully dissolve the PMMA layer. WSTs (Aquasol Corporation) were used as a stamp to pick up the MEAs. A 50 nm-thick transparent SiO_2_ was then deposited on the back of MEAs by electron beam evaporation. The WSTs and receiving PDMS substrate (Sylgard 184, Dow Corning, crosslinker:PDMS base = 1:10) were treated in a high power oxygen plasma cleaner (PDC-001-HP, Harrick Plasma) at 45 W for 2 minutes, attached together tightly, and thermally heated at 150 °C for 20 minutes. Immersing the samples in water removed the WSTs and released the MEAs.

#### Optical Measurement

A spectrophotometer (V-770 UV-vis/NIR, Jasco Inc.) measured the optical transmission spectra of the Ag NWs and Au-Ag NWs films.

#### Morphology Characterization

The morphology of the Au-Ag NWs networks was measured by SEM (PIONEER EBL, Raith Inc.) and TEM (Talos F200X G2 TEM, ThermoFisher Scientific).

#### Electrical Measurement

Sheet resistance was measured with a four-point probe (SRM-232, Guardian Manufacturing Inc.).

#### Electrochemical Measurement

EIS measurements were conducted with a Gamry potentiostat (Reference 600+, Gamry Instruments Inc.) at a frequency range from 1 Hz to 100 kHz in a 1 × PBS solution (Sigma-Aldrich). The measurements used a three-electrode configuration where the NWs microelectrodes, an Ag/AgCl electrode, and a platinum electrode served as the working, reference, and counter electrodes, respectively. For benchtop measurements, the sine wave signal (10 Hz, 20 mV peak-to-peak amplitude) was delivered by a PowerLab data acquisition system (PowerLab 16/35, ADInstruments Inc.) through a platinum electrode into a 1 × PBS solution (Sigma-Aldrich). The PSD signals were processed in MATLAB.

#### Mechanical Measurement

Mechanical performance of the Au-Ag NWs MEAs was assessed by a motorized test stand (ESM 1500, Mark-10). Electrochemical measurements were conducted after stretching.

#### FEA Simulations

Commercial FEA software Abaqus/CAE 2018 served as the tool to simulate the stretching of the Au-Ag NWs MEA structures. The device was clamped at the top and pulled unidirectionally from the bottom. The device was stretched up to 40% of the initial length. Shell elements with respective materials and thickness values were used for the proper representation of the device. The nonlinear static general procedure was carried out using Abaqus/Standard solver. The mesh included linear quadrilateral (S4) and linear triangular (S3) elements. The FEA model included 520953 elements and 535443 nodes in total, which were determined by a mesh sensitivity study. The analysis was performed on a high-end Dell server with 32 cores (Intel Xeon Gold 6144), 1 NVIDIA Tesla P100 GPU, and 384 GB RAM for fast simulations.

#### Animal Experiments

All animal procedures were approved by The George Washington University Animal Care and Use Committee and in accordance with recommendations from the panel of Euthanasia of the American Veterinary Medical Association and the National Institutes of Health Guide for the Care and Use of Laboratory Animals.

Adult male Sprague-Dawley rats were anesthetized by inhalation of 5% isoflurane vapor and humanely sacrificed by rapid excision of the hearts after toe pinch verification of cessation of all pain reflexes. Excised hearts were cannulated via the aorta, washed of blood, and retrograde perfused at constant pressure with a modified Tyrode’s solution (pH 7.4 at 37 °C, in mM: NaCl 118, KCl 4.7, MgSO_4_ 0.57, CaCl_2_ 1.25, KH_2_PO_4_ 1.17, NaHCO_3_ 25, Glucose 6). Aortic pressure was maintained from 70-80 mmHg. The hearts were allowed to stabilize for at least 10 minutes. The Au-Ag NWs MEAs were placed on the curvilinear epicardial surface of the left ventricle for electrical recording. The recorded EG signals were acquired using a 16-channel PowerLab data acquisition system (ADInstruments Inc.) at a sample frequency at 20 kHz and analyzed using LabChart software (ADInstruments Inc.). ECG signals were recorded using needle electrodes (MLA1203, ADInstruments Inc.) placed within the superfusion fluid that bathed the heart. After staining the heart with the potentiometric dye RH237 (Invitrogen™), the dye was energized using two 520 nm ultra-high power LEDs (Prizmatix). Optical action potentials were imaged with a CCD camera (iXon DV860, Andor Technology) at a frame rate of 490 frames per second, as previously described.^[44-45]^ Optical mapping data were analyzed using custom MATLAB algorithms.^[45]^ For electrical pacing, a platinum pacing electrode was placed at the base of the left ventricle and connected to a constant current source (Stimulus Isolator A385, World Precision Instruments) to deliver 3.6 mA of current with 5 ms pulse duration.

## Acknowledgements

Z. C., K. N., G. K., and X. S. contributed equally to this work. We thank The George Washington University Nanofabrication and Imaging Center for its facilities and device fabrication support. L. L. acknowledges the support of the National Science Foundation grants (ECCS 2011093 and CBET 2131682) and The George Washington University Cross-Disciplinary Research Fund. M. W. K. acknowledges the support of National Institutes of Health grants (HL146169 and HL147279). X. N. acknowledges the support of the National Science Foundation grant (ECCS 2030579).

