## Supplemental Figures 1-3 for "Transparent and stretchable metal nanowire composite recording microelectrode arrays"

### Supplementary Material

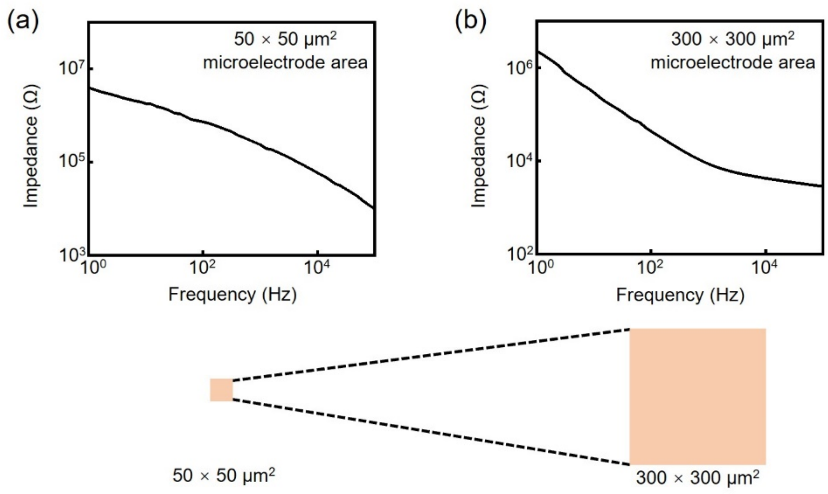


**Figure S1.** Impedance plots of a 50 × 50 $\mu m^2$ cellular scale Au-Ag NWs microelectrode (a) and a 300 × 300 $\mu m^2$ Au-Ag NWs microelectrode (b).

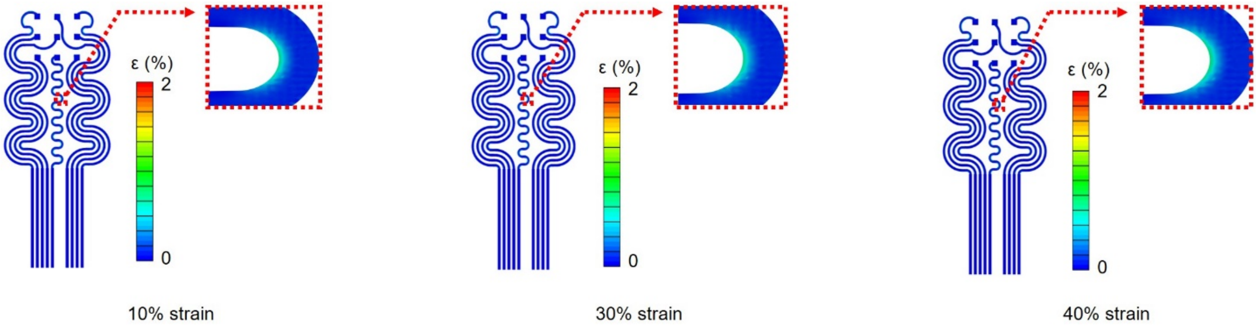


**Figure S2.** FEA strain distribution results of Au-Ag NWs MEAs and interconnects under 10% (left), 30% (middle), and 40% (right) uniaxial stretching.

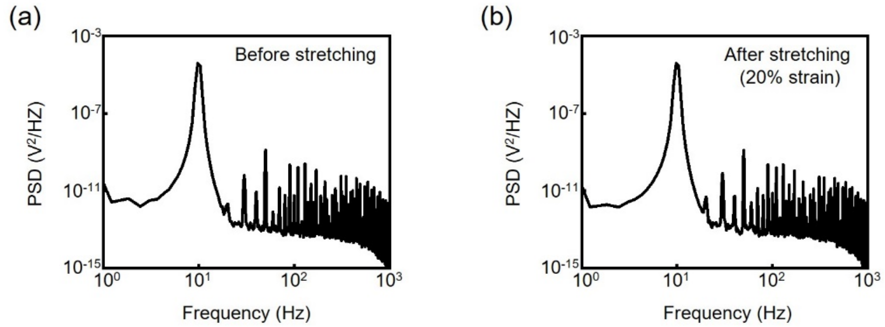


**Figure S3.** PSD of the electrical signals recorded by the pristine (a) and stretched (b) Au-Ag NWs microelectrodes.
